# Discovery and Characterization of Novel Lignocellulose-Degrading Enzymes from the Porcupine Microbiome by Synthetic Metagenomics

**DOI:** 10.1101/288985

**Authors:** Mackenzie Thornbury, Jacob Sicheri, Patrick Slaine, Landon J. Getz, Emma Finlayson-Trick, Jamie Cook, Caroline Guinard, Nicholas Boudreau, David Jakeman, John Rohde, Craig McCormick

## Abstract

Plant cell walls are composed of cellulose, hemicellulose, and lignin, collectively known as lignocellulose. Microorganisms degrade lignocellulose to liberate sugars to meet metabolic demands. Using a metagenomic sequencing approach, we previously demonstrated that the microbiome of the North American porcupine *(Erethizon dorsatum)* is replete with lignocellulose-degrading enzymes. Here, we report the identification, synthesis and partial characterization of four novel genes from the porcupine microbiome encoding putative lignocellulose-degrading enzymes; β-glucosidase, β-L-arabinofuranosidase, β-xylosidase, and an endo-1,4-β-xylanase. These genes were identified via conserved catalytic domains associated with cellulose- and hemicellulose-degradation, and phylogenetic trees were created to depict relatedness to known enzymes. The candidate synthesized genes were cloned into the pET26b(+) plasmid to enable inducible expression in *Escherichia coli (E. coli)*. Each candidate gene was cloned as a fusion protein bearing an amino-terminal PelB motif required for periplasmic localization and subsequent secretion, and a carboxy-terminal hexahistidine (6xHIS) tag to enable affinity purification. We demonstrated IPTG-inducible accumulation of all four fusion proteins. The putative β-glucosidase fusion protein was efficiently secreted but did not permit *E. coli* to use cellobiose as a sole carbon source, nor did the affinity purified enzyme cleave *p*-Nitrophenyl β-D-glucopyranoside (*p*-NPG) substrate *in vitro* over a range of physiological pH levels (pH 5-7). By contrast, the affinity purified putative endo-1,4-β-xylanase protein cleaved a 6-chloro-4-methylumbelliferyl xylobioside substrate over this same pH range, with maximal activity at pH 7. At this optimal pH, *K_M_*, *V*_max_, and *k_cat_* were determined to be 32.005 ± 4.72 μΜ, 1.16×10^-5^ ± 3.55×10^-7^ M/s, and 94.72 s^-1^, respectively. Thus, our synthetic metagenomic pipeline enabled successful identification and characterization of a novel hemicellulose-degrading enzyme from the porcupine microbiome. Progress towards the goal of introducing a complete lignocellulose-degradation pathway into *E. coli* will be accelerated by combining synthetic metagenomic approaches with functional metagenomic library screening, which can identify novel enzymes unrelated to those found in available databases.

**Author summary:** Plants depend on a mixture of complex polysaccharides including cellulose, hemicellulose and lignin to provide structural support. Microorganisms use enzymes to break down these polysaccharides into simple sugars like glucose that they burn for energy. These microbial enzymes have the potential to be repurposed for industrial biofuel production. Previously, we showed that bacteria in the porcupine gut produce enzymes that break down cellulose and hemicellulose. Here we report the identification and synthesis of four genes from the porcupine microbiome that encode enzymes that can break down complex plant polysaccharides, including a β-glucosidase, an β-L-arabinofuranosidase, a β-xylosidase, and an endo-1,4-β-xylanase. These genes were introduced into the model bacterium *Escherichia coli* (*E coli)* to test the properties of their respective gene products. All four enzymes accumulated following the induction of gene expression, but only the putative cellulose-degrading β-glucosidase and hemicellulose-degrading β-xylosidase and β-L-arabinofuranosidase enzymes were efficiently secreted out of the bacteria where they could access their target polysaccharides. Expression of the putative β-glucosidase did not allow *E. coli* to grow on cellobiose as a sole carbon source, and the purified enzyme failed to cleave *p*-NPG *in vitro*. By contrast, the purified putative endo-1,4-β-xylanase cleaved an artificial hemicellulose substrate *in vitro* across a range of acidic and neutral pH values, with maximal activity at pH 7. This study demonstrates the power of our approach to identify novel microbial genes that may be useful for industrial biofuel production.

## Introduction

The gut microbiome comprises thousands of microbial species encoding millions of genes that affect host physiology [1]. These microbes provide a genetic repository of digestive enzymes that convert complex lignocellulose polymers into simple fermentable sugars [2,3]. This repository of unique enzymes has the potential to be harnessed for industrial purposes [4]. Lignocellulosic biomass consists of cellulose, hemicellulose, and lignin, typically found in a 4:3:3 ratio [3]. Microbial enzymes have been shown to catalyze the degradation of cellulose and hemicellulose into monosaccharides that can be utilized for the production of ethanol biofuel [5,6,7,8]. However, hemicellulose has not yet been extensively investigated as a biofuel substrate. Moreover, lignin has a complex chemical structure that impedes chemical and enzymatic hydrolysis. Wi *et al*. recently demonstrated a new hydrogen peroxide pre-treatment that improves downstream biocatalytic hydrolysis of lignocellulose by removing lignin [9]. Overall, microbial lignocellulose-degrading enzymes remain a largely untapped resource for biofuel production.

The North American Porcupine, *Erethizon dorsatum*, is a hind-gut fermenter with an enlarged cecum packed with microbes that aid digestion of lignified plants, coniferous and deciduous cambium (inner bark), and flowers [10]. Using metagenomic sequencing, the 2016 Dalhousie international Genetically Engineered Machine (iGEM) team determined that the porcupine microbiome is replete with microbial enzymes with potential lignocellulose-degrading properties [11]. Furthermore, using shotgun metagenomic and 16S sequencing, the authors confirmed that host diet influences gut microbial diversity and metabolic function; they reported that herbivores like beavers and porcupines had elevated levels of cellulolytic genes in their microbiomes compared to carnivores like the Arctic wolf and coyote.

These findings inspired the 2017 Dalhousie iGEM team to continue studying enzymes from the porcupine microbiome with the long-term objective of engineering a novel lignocellulose-degradation pathway in *E. coli* for biofuel applications. We created our own synthetic metagenomic pipeline to identify and characterize putative cellulose- and hemicellulose-degrading enzymes from existing datasets.

## Results

### Identification and Cloning of Putative Microbial Enzymes via a Synthetic Metagenomic Pipeline

We used a metagenomic sequencing pipeline (Fig. 1A) to identify four microbial genes from porcupine fecal samples predicted to encode putative cellulose- or hemicellulose-degrading enzymes; a β-glucosidase, an β-L-arabinofuranosidase, a β-xylosidase, and an endo-1,4-β-xylanase (Figs 1B, 2A). These genes were identified by similarity of predicted primary amino acid sequences to conserved domains from known enzymes found in the Research Collaboratory for Structural Bioinformatics Protein Data Bank (Fig 1B). β-glucosidase enzymes catalyze the final step in cellulose degradation by converting cellobiose disaccharides to glucose monomers (Fig 2B), whereas β-L-arabinofuranosidase, β-xylosidase, and endo-1,4-β-xylanase catalyze sequential steps in the degradation of hemicellulose to xylose monomers (Fig. 2C). This collection of putative enzymes is insufficient to achieve full degradation of complex lignocellulose substrates, but their synthesis and characterization serves as an important test of our synthetic metagenomic pipeline.

**Fig 1.**
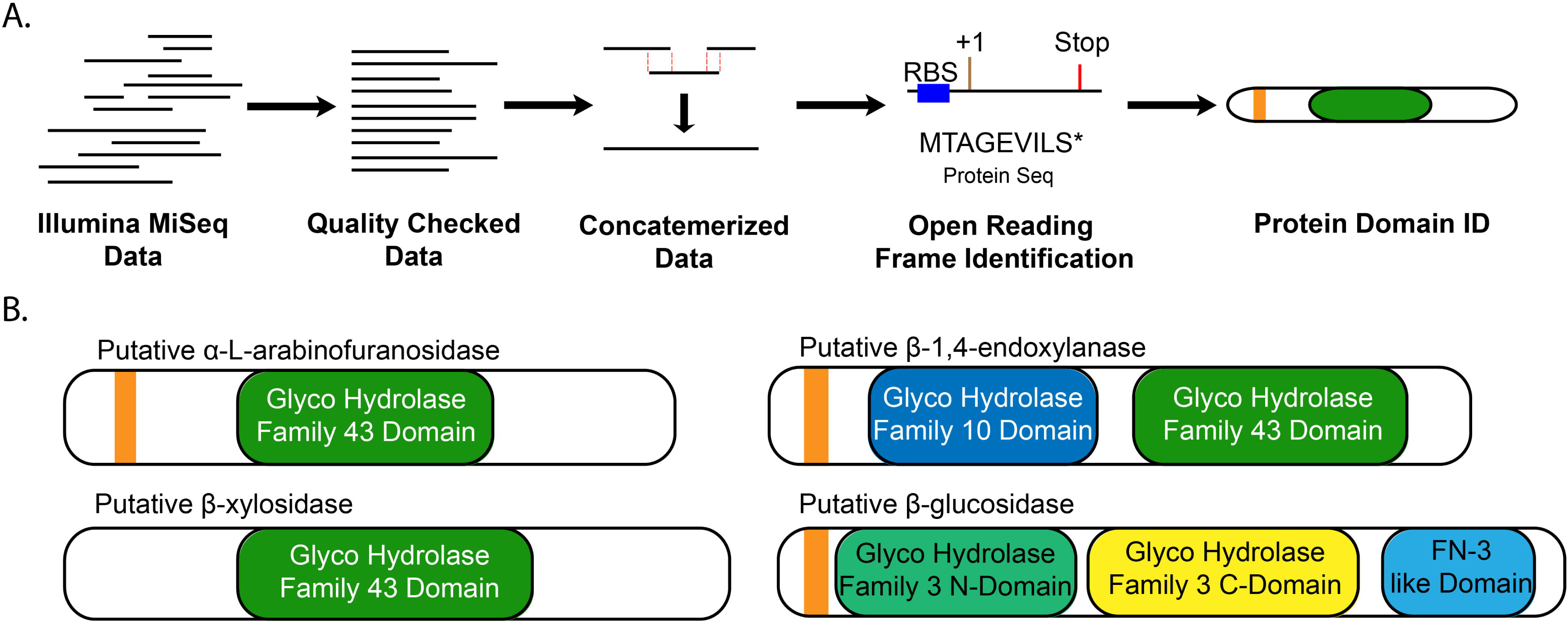
Identification of four genes from the porcupine microbiome with putative cellulose- and/or hemicellulose-degrading activity using a metagenomic sequencing pipeline. The bioinformatic pipeline (A) began with Illumina MiSeq data previously collected from a porcupine fecal DNA sequencing project [11]. Reads were checked for quality and trimmed, concatemerized via MegaHit [12], and open reading frames were identified using Prodigal [13]. Protein sequences of interest were identified by pHMMER [14] using various protein databases and were selected for matches of interest based on e-value selection. (B) Top putative microbial enzymes identified by the metagenomic sequencing pipeline; putative signal sequences are shown in orange and predicted conserved protein domains are shown.

**Fig 2.**
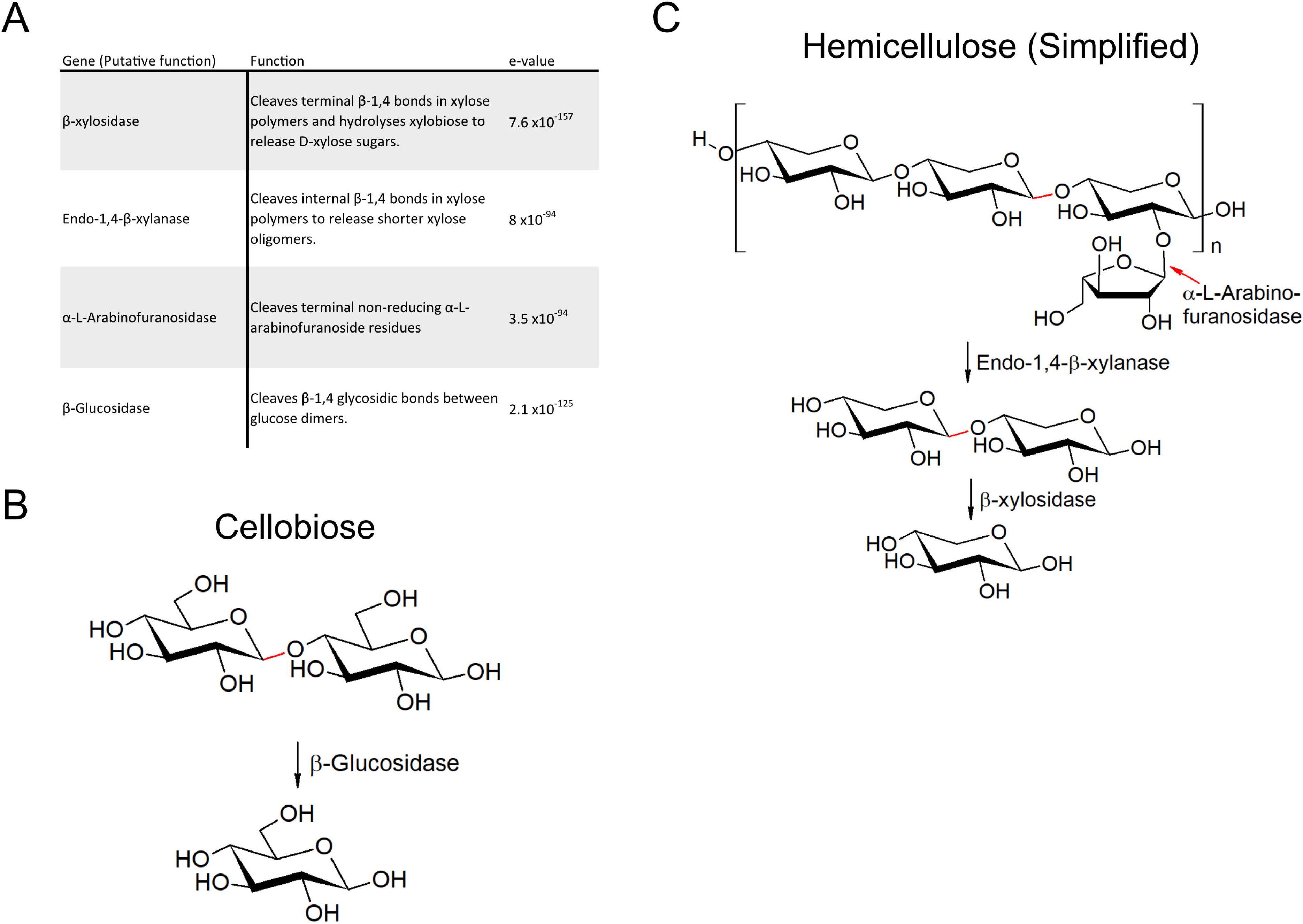
Four putative cellulose-/hemicellulose-degrading enzymes identified from the porcupine microbiome. A) Functional description of putative enzymes with corresponding e-values. The e-value is a measure of confidence, with lower values denoting higher confidence. B) The catabolic pathway that converts cellobiose to glucose C) The catabolic pathway that converts hemicellulose to xylose.

### *In Silico* Analysis of Putative Enzymes

The predicted primary amino acid sequence of each candidate gene was used to query the NCBI non-redundant protein sequence database using the Basic Local Alignment Search Tool (BLAST) [15]. Each of our four putative enzymes from the porcupine microbiome were most closely related to enzymes encoded by anaerobic bacteria. Specifically, our putative β-xylosidase was 75% identical (100% coverage) to a β-xylosidase from *Butyrivibrio* sp. CAG:318. Our putative β-glucosidase was 73% identical (99% coverage) to a β-glucosidase encoded by *Bacteroides faecis* MAJ27. Our putative β-L-arabinofuranosidase was 63% identical (92% coverage) to a glycosyl hydrolase (GH) family 43 protein encoded by *Prevotella* sp. CAG:732. The putative endo-1,4-β-xylanase was 55% identical (99% coverage) to a hypothetical protein BHV73_02415 encoded by *Bacteroides* sp. 44_46 and predicted to contain GH10 and GH43 domains.

Phylogenetic analysis of all four putative gene products was performed using the 22 closest homologs for each. The putative endo-1,4-β-xylanase and β-glucosidase enzymes clustered closely with their homologs in a clade (Fig 3). These clades were strongly supported by high bootstrap values (100 and 100, respectively) on each node, generated from compiling 100 separate trees. The putative β-L-arabinofuranosidase did not fall into a clade, but did cluster with its closest homologs, whereas β-xylosidase clustered poorly. Next, predicted amino acid sequences of all four putative enzymes were aligned with closest homologs to determine conservation of key catalytic residues (Figs S1A-S1D). A key aspartic acid is conserved in the catalytic domain of the putative β-glucosidase; this residue was conserved across all 22 proteins analyzed, with 4 examples aligned to the putative enzyme in Fig. S1A. By contrast, the catalytic domain of endo-1,4-β-xylanase consists of two conserved aspartic acids and a glutamic acid; all three residues were perfectly conserved over all 22 proteins analyzed, including the predicted protein, and 4 example sequences were aligned to the putative endo-1,4-β-xylanase in Fig. S1B. β-L-arabinofuranosidase similarly has two conserved aspartic acids and a single glutamic acid responsible for activity, that shows perfect conservation against all proteins analyzed (S1B). The putative β-xylosidase showed low conservation from amino acids 190–230, but the conserved catalytic domain requiring the two conserved aspartic acids and a single glutamic acid were also found to be conserved across all sequences analyzed. Identifying and confirming the catalytic sites provides evidence that these proteins may function in a biological system and require functional assays to determine activity by *de novo* synthesis of these open reading frames.

**Fig 3.**
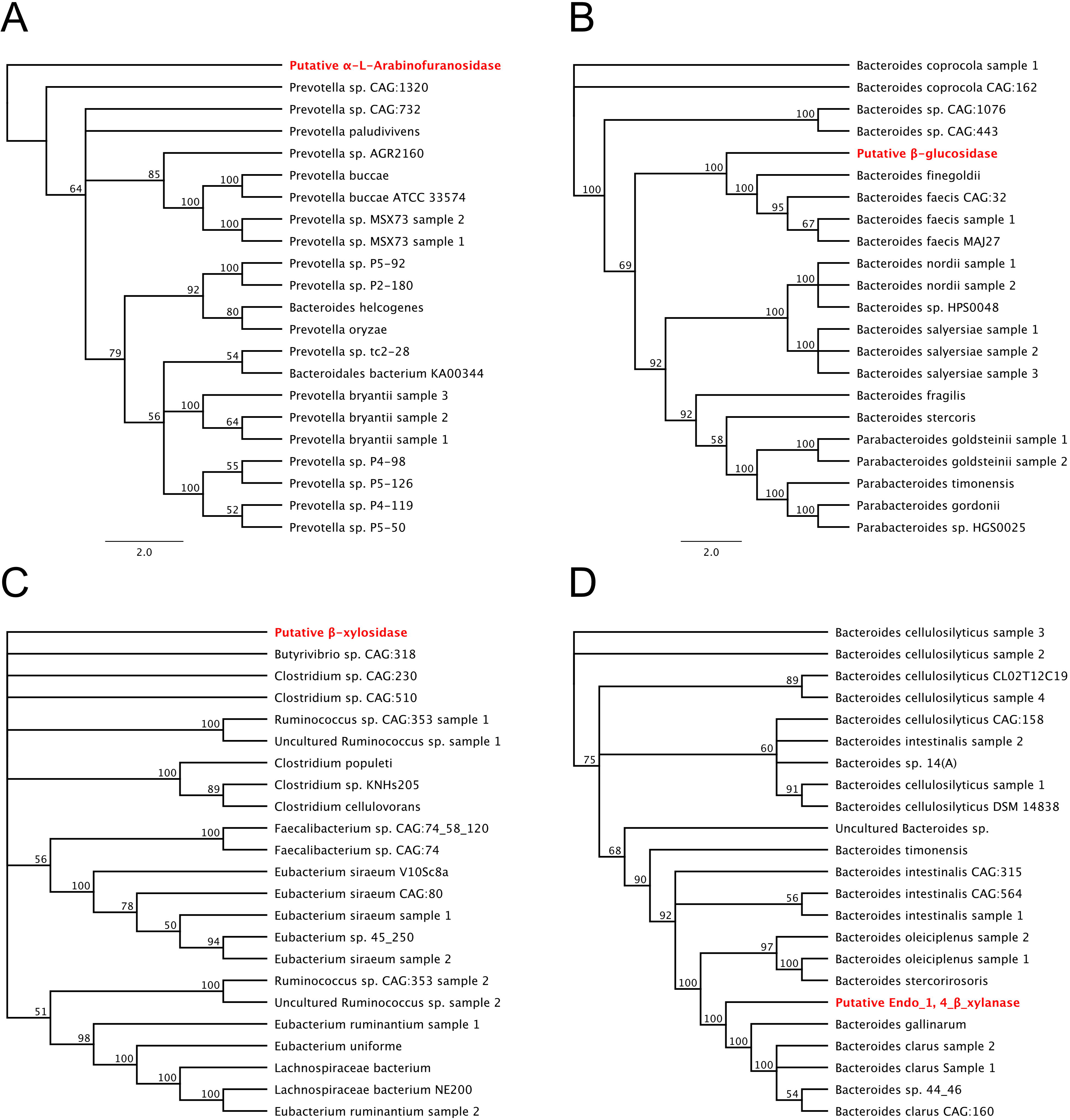
Phylogenetic analysis of putative enzymes. Phylogenetic analysis of A) β-L-arabinofuranosidase, B) β-glucosidase, C) β-xylosidase, and D) Endo-1,4-β-xylanase were created from primary amino acid sequence that were aligned using ClustalW and generated using RAxML 7.2.8. Sequences were taken from highly related proteins from the BLASTP database. Trees were replicated 100 times for statistical strength with bootstrap values presented at each node. Putative enzymes are highlighted in red.

### Synthesis, Expression and Secretion of Putative Enzymes in *E. coli*

The four putative microbial enzymes were synthesized and cloned into the pET26b(+) vector to enable IPTG-inducible gene expression in *E. coli*. Because directed secretion of enzymes provides access to extracellular lignocellulosic substrates, each candidate gene was cloned as a fusion protein bearing an amino-terminal PelB motif required for periplasmic localization and subsequent secretion. A hexahistidine (6xHIS) tag was added to carboxy-termini of each fusion protein to enable affinity purification. Log-phase *E. coli* cultures were treated with IPTG to induce transgene expression, followed by harvest of cell supernatant, periplasm and total cell fractions. Specifically, proteins in the supernatant were harvested by trichloroacetic acid (TCA) precipitation and periplasmic proteins were harvested by cold osmotic extraction as previously described [16]. These fractions were subjected to SDS-PAGE and immunoblotting to detect 6XHIS fusion proteins (Fig 4). All four putative enzymes accumulated in the total cell fraction: β-xylosidase at ~51 kDa, endo-1,4-β-xylanase at ~81 kDa, β-L-arabinofuranosidase at ~61 kDa, and β-glucosidase at ~84 kDa (Fig 4A). Among these putative enzymes, only the endo-1,4-β-xylanase failed to translocate to the periplasm, and it accumulated in the cell pellet fraction (Figs 4B, 4C).

**Fig 4.**
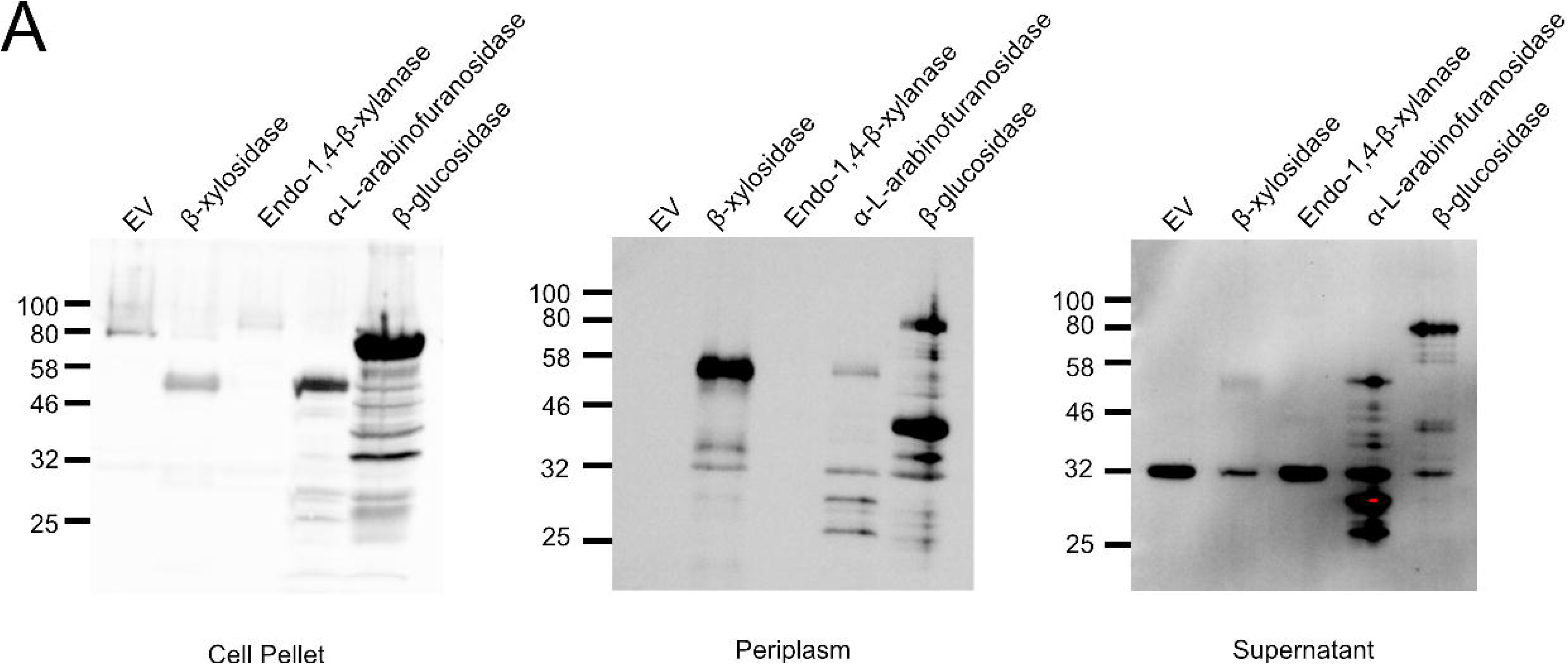
Secretion Assay of Putative Enzymes. Log-phase BL21(DE3) *E. coli* containing a putative enzyme or empty vector control were treated with 0.1 mM IPTG to induce T7-promoter-dependent transcription. After 3 hours, A) cells were lysed, B) cells were treated with cold osmotic shock to extract periplasmic protein fraction, or C) the supernatant was filtered, and protein was precipitated by addition of TCA. All protein fractions were prepared for SDS-PAGE and analyzed by western blot with an anti-His-antibody.

### Isolation and Characterization of a Putative β-glucosidase

Treatment of transformed log-phase *E. coli* cultures with IPTG caused accumulation of the known beta-glucosidase DesR [17,18] which we isolated by affinity purification and detected by immunoblotting with an anti-6XHIS antibody (Fig 5A). Our putative microbial β-glucosidase fusion protein also accumulated and was affinity purified (Fig 5B). Consistent with published reports, purified DesR efficiently cleaved a *p*-Nitrophenyl β-D-glucopyranoside (*p*-NPG) substrate *in vitro* at pH 7 (Fig 5C). By contrast, our β-glucosidase failed to cleave *p*-NPG over a broad pH range (Fig 5D). Moreover, expression of the putative β-glucosidase in *E. coli* did not enable growth on cellobiose as a sole carbon source. Taken together, these findings indicate that our putative β-glucosidase does not function in conventional β-glucosidase assays.

**Fig 5.**
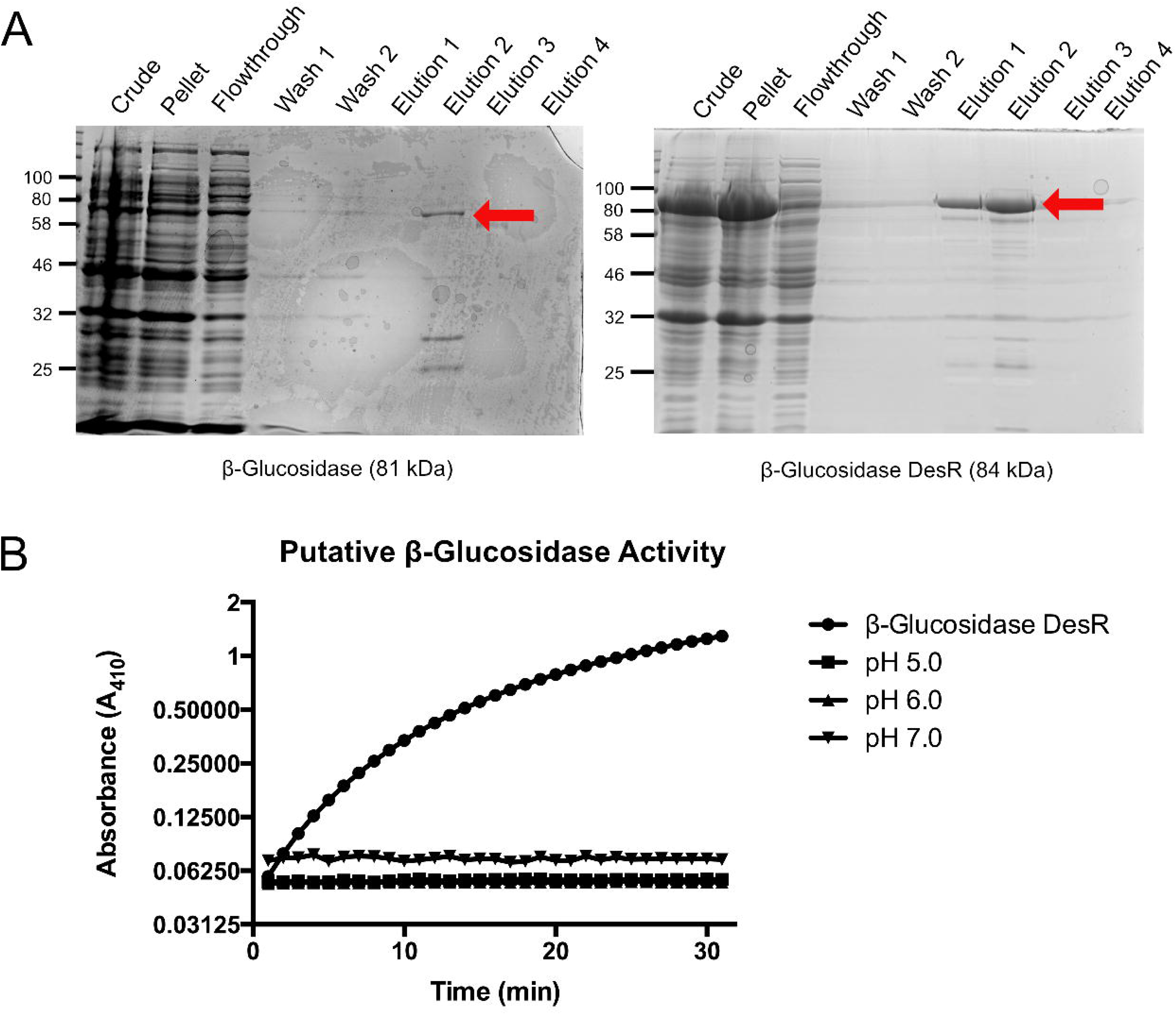
Purification and Characterization of β-glucosidase. A) Putative β-glucosidase and positive control β-glucosidase DesR were purified by 6x His purification. Red arrow indicates relevant band B) Activity of putative β-glucosidase against *p*-NPG at pH 5.0, 6.0 and 7.0, plotted with positive control; β-glucosidase DesR at pH 7.0.

### Isolation and Characterization of a Putative Endo-1,4-β-xylanase

We successfully purified our putative endo-1,4-β-xylanase using the protein production and isolation protocol described above (Fig 6A). Enzyme activity was assessed using 6-chloro-4-methylumbelliferyl xylobioside (CMU-X_2_) substrate and a modified protocol developed by Hallam and Withers [19]. When cleaved by an active enzyme, CMU is released from xylobiose, causing fluorescence emission. CMU-X_2_ (100 μM) was combined with 0.6 μg of purified putative endo-1,4-β-xylanase over a wide pH range. Immediately upon addition of enzyme, the fluorophore was excited at 365 nm and emission was read at 450 nm; one measurement was taken every minute for 30 minutes. The increase in raw fluorescence units (RFUs) over time indicated cleavage of the CMU-X_2_ substrate by the putative endo-1,4-β-xylanase; this reaction was catalyzed most efficiently at pH 7 (Fig 6B). The DesR β-glucosidase served as a negative control in these assays and did not cleave the CMU-X_2_ substrate (Figure S3).

To investigate the thermostability of the putative endo-1,4-β-xylanase, we incubated samples containing 0.6 μg of purified enzyme for 30 minutes at 37°C, 50°C, 60°C, or 70°C before conducting an *in vitro* CMU-X_2_ cleavage assay as described above. Enzyme activity was abolished after 30-minute incubation at any of these temperatures (Fig 6D). A Michaelis-Menten plot was generated for endo-1,4-β-xylanase activity over a range of CMU-X_2_ concentrations (Fig 6E). From this, kinetic constants for endo-1,4-β-xylanase were calculated using a Lineweaver-Burk double reciprocal plot (Fig 6F). Endo-1,4-β-xylanase had a calculated *K_M_* value of 32.005 ± 4.72 μM for CMU-X_2_ and a V_max_ of 1.16×10^-5^ ± 3.55×10^-7^ M/s. The calculated turnover number (*k*_cat_) for endo-1,4-β-xylanase was found to be 94.7 s^-1^. Finally, *k*_cat_/*K_M_* was calculated to be 2.96×10^6^ M^-1^s^-1^.

**Fig 6.**
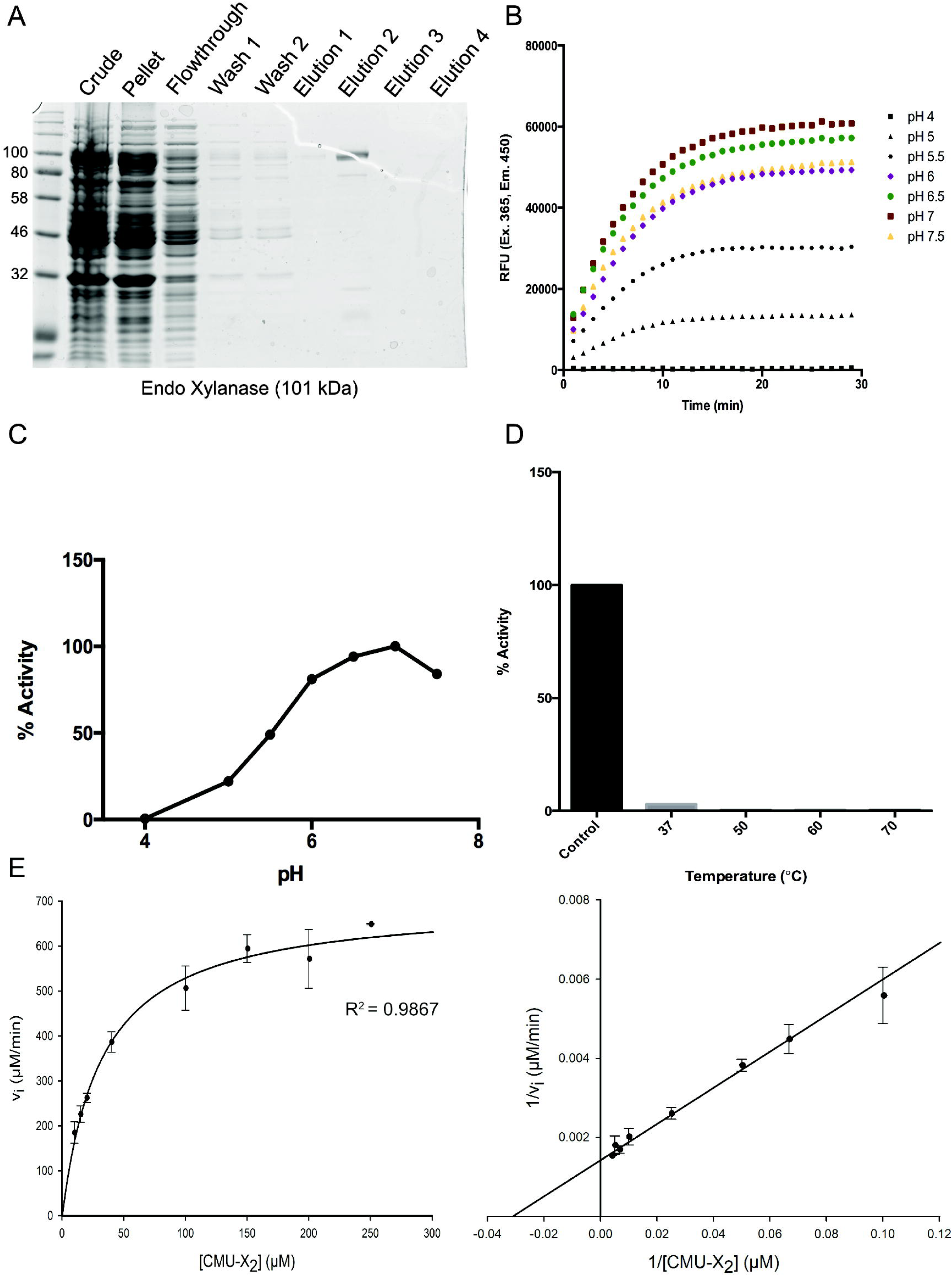
Characterization of Endo-1,4-β-xylanase. A) Putative endo-1,4-β-xylanase was purified by 6x His purification. B) 0.6 μg of endo-1,4-β-xylanase was combined with 100 μM of CMU-X_2_ to assess enzyme activity at different pH. Samples were assessed via fluorescence in sodium citrate buffer (pH 4, 5, 5.5, 6) or sodium phosphate buffer (pH 6.5, 7, 7.5). A reading of raw fluorescent units (450 nm) was taken every minute for 30 minutes at 37°. C) Relative activity of endo-1,4-β-xylanase at different pH compared to pH 7. D) Thermostability of endo-1,4-β-xylanase was assessed. Samples containing 0.6 μg of endo-1,4-β-xylanase in 50 mM sodium phosphate buffer were incubated for 30 minutes at 37°, 50°, 60°, or 70° before being introduced to 100 μΜ of CMU-X_2_ and read as before. Control is unincubated endo-1,4-β-xylanase. E) Michaelis-Menten plot was generated with initial reaction rate against substrate concentrations above at 37°, pH 7. F) Lineweaver-Burk plot was used to generate kinetic constants *K*_M_, V_max_, and *k*_cat_.

## Discussion

The Dalhousie 2016 iGEM team previously demonstrated that the porcupine microbiome is a particularly rich source of lignocellulose-degrading enzymes [11]. We developed a synthetic metagenomic pipeline to mine the porcupine microbiome for novel enzymes with useful properties. We used stringent selection criteria to maximize our chances of discovering *bona fide* cellulose/hemicellulose-degrading enzymes. We identified and synthesized four previously uncharacterized genes encoding putative cellulose- and hemicellulose-degrading enzymes from the porcupine microbiome: a β-glucosidase, an β-L-arabinofuranosidase, a β-xylosidase, and an endo-1,4-β-xylanase. We thoroughly analyzed these putative enzymes by phylogenetic analysis and identified key conserved residues in proposed catalytic sites. These putative enzymes were affinity purified, and we demonstrated clear *in vitro* activity of a purified putative endo-1,4-β-xylanase, with optimal activity at pH 7, consistent with the neutral pH of the porcupine cecum [20]. This marks the first discovery of a functional hemicellulose-degrading enzyme from the porcupine microbiome, and it demonstrates the power of our synthetic metagenomic approach.

The microbial genome that encodes our novel endo-1,4-β-xylanase is unknown, but it has a Shine-Dalgarno (SD) Sequence found only in bacterial genes, and phylogenetic analysis revealed that it is most closely related to uncharacterized proteins with glycosyl hydrolase (GH) family 10 and 43 domains from *Bacteroides sp* as described by the Carbohydrate Active Enzymes Database (CAZy, http://www.cazy.org/) [21]. GH10 domains have been linked to endo-1,4-β-xylanase, endo-1,3-β-xylanase, tomatinase and/or xylan endotransglycosylase activity, while GH43 domains are commonly found in β-xylosidase and/or β-L-arabinofuranosidase enzymes [21]. Because our candidate endo-1,4-β-xylanase cleaved a modified xylobiose substrate *in vitro*, we inferred that the GH10 domain was functional in the context of the affinity-purified fusion protein, despite the addition of an amino-terminal PelB domain and carboxy-terminal 6XHis affinity purification tag. This enzyme displayed activity across a wide range of pH values (pH 4-7.5), but lacked thermostability, quickly losing activity after 30 min incubation at a range of temperatures as low as 37°, despite the fact that the porcupine deep core body temperature is approximately 37° [20,22]. This thermo-instability makes our novel endo-1,4-β-xylanase less useful for industrial applications compared to known thermophilic endo-1,4-β-xylanases [23]. Enzyme thermostability can be improved via directed evolution; Stephens *et. al*. used error-prone PCR to increase endo-1,4-β-xylanase thermostability, creating a superior variant with a single amino acid substitution [24]. Advances in computational prediction and protein folding models can also inform rational mutagenesis strategies to increase enzyme thermostability [25, 26, 27]. Such approaches may be used to enhance the thermostability of our novel endo-1,4-β-xylanase. We calculated a turnover rate (*k*_cat_) of 94.7 s^-1^ for our novel endo-1,4-β-xylanase, which compares favorably with other known enzymes. Using beechwood xylan as a substrate, He *et. al*. reported that an endo-1,4-β-xylanase isolated from the fungus *Trichoderma reesei* had a relatively high *k_cat_* of 139.7 s^-1^[28]. Using the same substrate, Xu *et. al*. reported a lower rate of turnover of 47.34 s^-1^ for a microbial xylanase containing a GH10 family domain isolated from the feces of the black snub-nosed monkey *(Rhinopithecus bieti)* [29]. A GH11 family endo-1,4-β-xylanase isolated from the fungus *Fusarium oxysporum* also had a low rate of turnover (0.27 s^-1^) of RBB-xylan substrate, a chromogenic derivative of beechwood xylan [30]. Like other kinetic constants, turnover rates are substrate-specific. Because we were the first to use CMU-X_2_ to calculate kinetic constants for an endo-1,4-β-xylanase, we cannot directly compare our findings to studies that employ beechwood xylan or RBB-xylan. Future studies should characterize kinetic constants of our endo-1,4-β-xylanase on more conventional substrates such as beechwood, birchwood, and oat-spelt xylan.

Among the other three putative enzymes identified in this study, the β-glucosidase accumulated to high levels, translocated to the periplasm and was efficiently secreted, yet it lacked *in vitro* activity. The putative β-L-arabinofuranosidase was also efficiently secreted, but its enzymatic activity remains to be tested as we seek out appropriate substrates. By contrast, β-xylosidase was difficult to purify and was often retained in the insoluble pellet fraction. Insolubility can occur when a protein aggregates before it can fold properly [31]. To promote proper folding and solubility of our putative β-xylosidase, we reduced IPTG levels from 1.0 mM to 0.1 mM and reduced the culture temperature to 20 °C during the time of induction. Despite these precautions, the amount of affinity purified β-xylosidase remained insufficient for subsequent *in vitro* enzyme assays. It is possible that *E. coli* lacks appropriate chaperone activity required to efficiently fold this enzyme.

Our synthetic metagenomic pipeline is a powerful tool for discovery, but because it infers functional relationships from homology to previously characterized sequences in a database, it will likely fail to identify greatly divergent proteins with desirable properties. It may also identify putative enzymes that appear to have conserved functional domains, but lack the function predicted by homology. By contrast, functional metagenomic library screens rely on functional assays for gene discovery. Thus, functional metagenomics provides a convenient approach to new gene discovery that nicely complements sequence-based approaches, but with greater potential for discovery of truly novel genes that don’t resemble those in existing databases. Recently, using functional metagenomics, Cheng, *et. al*. discovered three novel β-galactosidase enzymes, two of which had conserved domains, and one of which was part of a previously undiscovered enzyme family [32]. To complement our synthetic metagenomics approach, we plan to create and screen a functional metagenomic library from porcupine microbiome DNA to discover novel lignocellulose-degrading enzymes.

## Materials and Methods

### Identification of Open Reading Frames

Metagenomic analysis of Illumina MiSeq data was conducted using our previously published protocols [11]. FASTQC and BowTie2 were used to inspect reads for overall quality and contaminants from sequencing. Reads were trimmed to 400 bp in length to remove low-quality terminal sequences from further analysis. MegaHIT alignment software processed reads in FASTq format and stitched reads into longer contigs by identifying overlapping coding regions [12]. Prodigal was used to identify open reading frames (ORFs) by searching sequences in six frames across both DNA strands [13]. A ‘-c’ command modifier in Prodigal was used to ensure the program only detected ORFs with both start and stop codons present. Prodigal was instructed to search for Shine-Dalgarno sequences required for ribosome binding to prokaryotic mRNAs, and non-canonical start codons CUG, GUG and ACG which are typically found in up to 10% of prokaryotic ORFs; these products are often overlooked by conventional searches and absent from many databases [33,34]. These restrictions were expected to limit our hits to prokaryotic genes, rather than genes from eukaryotes like fungi and protists that lack Shine-Dalgarno sequences.

### In Silico Protein Function Predictions

pHmmer was used to identify putative function of protein domains [14,35]. Protein domains and possible functions were identified using the Research Collaboratory for Structural Bioinformatics Protein Data Bank [36]. e-values were calculated to compare domains identified in putative proteins to known domains in the database [36]. Putative proteins with the lowest e-values were queried against the Basic Local Alignment Search Tool (BLAST) database using pHmmer to identify proteins with major protein domain conservation. Selected candidate genes were codon-optimized for *E. coli* and synthesized by Integrated DNA Technologies (IDT, Coralville, IA, USA) as gBlock gene fragments.

### Amino acid sequence analysis

Phylogenetic trees were generated and interpreted in Geneious R 8.1.8. using RAxML version 7.2.8 with the protein model GAMMA LG (algorithm: Rapid bootstrapping with 100 replicate trees for statistical power) [37].

Amino acid alignments were completed using ClustalW alignment with the cost matrix: BLOSUM with a gap open cost of 10 and a gap extend cost of 0.1 [38].

Putative genes were submitted to GenBank with the accession numbers as follows: β-Glucosidase MH590637, α-L-arabinofuranosidase MH590638, β-xylosidase MH590639, Endo-1,4-β-xylanase MH590640.

### Gene Cloning

Candidate genes were PCR-amplified from IDT gBlock gene fragments with Phusion High-Fidelity DNA Polymerase according to manufacturer’s instructions (New England Biolabs (NEB), Ipswich, MA, USA). PCR products were purified using the QIAquick gel extraction kit protocol (Qiagen Inc., Toronto, ON, Canada) (Table 1) and cloned into the pET26b(+) expression plasmid (MilliporeSigma); this plasmid enables the creation of fusion proteins with PelB leader sequences required for translocation to the periplasm after which they can be secreted into the extracellular space [39]. Thus, by fusing our putative enzymes to PelB we increased the likelihood of secretion to the extracellular space to access lignocellulosic substrates. Candidate genes and pET26b(+) were digested with restriction endonucleases (NEB) indicated in Table 1 for 1 hour at 37°. Digested DNA was subjected to agarose gel electrophoresis on a 0.8 % agarose gel and purified using the QIAquick gel extraction kit according to manufacturer’s instructions (Qiagen Inc.), then ligated with pET26b(+) plasmid DNA using T4 DNA ligase (NEB). Ligation products were transformed into chemically competent Stbl3 *E. coli* via heat-shock transformation [40]. Specifically, 5 μL of ligation products were added to 50 μL of *E. coli* suspension in Luria-Bertani (LB) broth, and following heat-shock transformation, 250 μL of LB broth was added during the 1 hour recovery stage and the mixture was subsequently plated on LB agar + 25 μg/ml kanamycin. Plates were incubated at 37 °C for 18–24 hours to allow the growth of transformants. Colonies were picked and inoculated into 5 mL of LB broth, grown overnight to saturation, and plasmid DNA was extracted via QIAprep Spin Miniprep Kit (Qiagen, Inc.). Plasmids were screened by restriction digestion, and processed for Sanger sequencing (Genewiz, South Plainfield, NJ, USA).

**Table 1.**
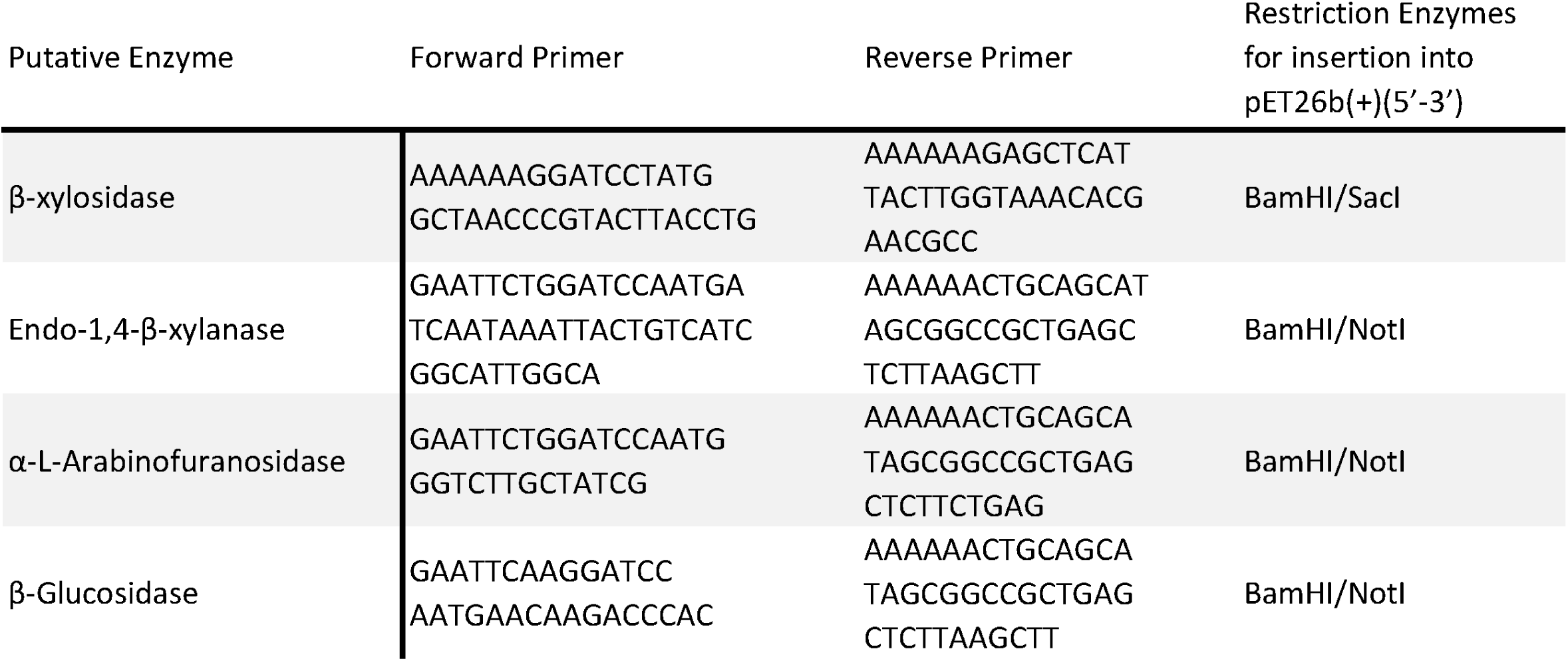
PCR Primers for amplification from Gene Blocks

The pET26b(+) vector contains a 6xHIS tag C terminal of the multiple cloning site (MCS). Initially, the candidate genes were cloned with a stop codon interrupting the 6xHIS tag. Site directed mutagenesis was used to delete the stop codon, restoring the C-terminal tag. The constructs were amplified using Phusion High-Fidelity DNA Polymerase according to manufacturer’s instructions (NEB). A small portion of PCR product was ran on a 0.8% agarose gel to confirm size. Once confirmed for proper size, the PCR product was purified with QIAquick PCR Purification Kit according to manufacturer’s instructions (Qiagen). After purification, the 6xHIS tagged constructs were ligated using T4 DNA ligase for 1 hr at 37 °C (NEB).

### Inducible Protein Expression

All pET26b(+) 6xHIS tagged candidate genes were transformed into BL21(DE3) *E. coli* to enable inducible protein expression. Selected colonies were inoculated into 5 mL of LB broth and incubated at 37°C in a shaker (220 RPM) overnight. The overnight culture was diluted 1:100 in 2.5 mL LB broth and incubated in a shaker until the OD_600_ of the culture reached 0.5–0.8. Once in log phase, 0.1 mM IPTG (Thermo Fisher Scientific, Waltham, MA, USA) was added to induce protein expression. After 3 hours of shaking incubation at 37 °C, bacteria were spun at 16000 xg for 2 min, supernatant was collected, and pellets were harvested in 2x electrophoresis sample buffer (ESB) or subjected to periplasmic fractionation. Protein was collected from the supernatant as previously described by Sarty *et al*. with some modification [41]. Specifically, 2.5 mL of clarified supernatant was passed through 0.2 μm PEL syringe filter and place on ice for 10 min before trichloroacetic acid precipitation. The periplasmic fraction was isolated using previously described cold osmotic shock [16]. Briefly, after the bacteria were pelleted, supernatant was removed, and the pellet was resuspended in 625 μL Cell Fractionation Buffer 1 (0.2 M Tris-HCl (pH 8.0), 200 g/L sucrose, 0.1 M EDTA) then incubated at 4 °C for 20 min with regular inversion. After incubation, the suspension was centrifuged at 16000 xg for 15 min at 4 °C and supernatant was discarded. The pellet was resuspended in 625 μL Cell Fractionation Buffer 2 (0.1 M Tris-HCl (pH 8.0), 0.005 M MgSO4, 0.2% SDS, 1% Triton X-100) and incubated for 20 min at 4 °C with regular inversion. The suspension was centrifuged as before, and the supernatant was collected as the periplasmic fraction. TCA precipitation was done to concentrate the protein. The vector pET26b(+) (EV) was used as a negative control. Proteins were separated by SDS-PAGE and transferred to PDVF and for western blot analysis using mouse penta HIS antibody (Qiagen, Cat. 34660).

### Protein Purification

Recombinant strains pET26b(+)-endo-1,4-β-xylanase and pET26b(+)-β-glucosidase were grown in 5 mL LB broth with 25 μg/mL kanamycin from 6 hrs at 37 °C, shaking, from a single clone. After initial incubation, 1 mL of culture was used to inoculate 100 mL of fresh LB broth with 25μg/mL kanamycin and 1mM IPTG for overnight induction at 30 °C, shaking. After induction, the cultures were centrifuged at 3220 xg for 20 min and supernatant was decanted from pellet. The cell pellets were resuspended in Wash Buffer (20 mM Na_2_PO_4_, 500 mM NaCl, pH 8.0) and sonicated for a total of 6 min in 30 sec intervals on ice, then centrifuged at 8000 xg for 30 min at 4 °C. Subsequently, supernatant was loaded into a column containing HisPur Cobalt Resin (Thermofisher) pre-equilibrated with Wash Buffer. The column was washed with 6 volumes of Wash Buffer, then protein was eluted in Elution Buffer (150 mM imidazole, 20 mM Na_2_PO_4_, 500 mM NaCl, pH 8.0). Glycerol was added to the purified protein at a final concentration of 10% to maintain protein folding.

Recombinant strain pET26b(+)-β-xylosidase was difficult to purify and a specialized protocol was used as previously described by Zimmermann *et al* [42]. Briefly, 1L of LB broth (no antibiotic) was inoculated with 5 mL of overnight pET26b(+)-β-xylosidase culture (containing kanamycin). The culture was shaken at 37 °C at 220 rpm until the OD_600_ was between 0.4 and 0.7. Once an appropriate density was reached, the culture was incubated at 42 °C for 10 min, recovered at 37 °C for 20 min, placed on ice for 30 min, and recovered again at 37 °C for 30 min. Once heat- and cold-shocked, the culture was induced with 0.1 mM IPTG and incubated overnight at 20 °C shaking. The following morning the culture was centrifuged and purified as described above.

### *In vitro* Activity Assays

The activity of β-xylosidase and β-glucosidase was tested using *p*-nitrophenol (*p*NP) derivatives: *p*NP-β-d-xylopyranoside and *p*NP-β-d-glucopyranoside, respectively. When cleaved by active enzyme, these compounds release *p*NP which can be measured by absorbance at 410 nm. General experimental procedure combined 900 μL of 5 mM *p*NP derivative in appropriate buffer, which was used to blank the spectrophotometer (Nanodrop One, Thermo Scientific), once 100 μL of appropriately diluted enzyme was added, the absorbance was measured at 410 nm every minute for 30 min. Optimum pH was determined by using three different buffers (50 mM) at pH 5: citrate buffer, pH 6: phosphate buffer, and pH 7: phosphate buffer.

The enzymatic activity of endo-1,4-β-xylanase was determined using 6-chloro-4-methylumbelliferyl xylobioside (CMU-X_2_). When cleaved by an active enzyme, 6-chloro-4-methylumbelliferyl is released from xylobiose and fluoresces at 450 nm. One hundred micromolar of CMU-X_2_ was added to 0.6 μg of HIS-purified endo-1,4-β-xylanase in 50 mM sodium citrate buffer (pH 4, 5, 5.5, 6) or sodium phosphate buffer (pH 6.5, 7, 7.5) in a 96-well plate (Grenier). Fluorescence was measured (Ex. 365, Em. 450) at 37° using a plate reader (Molecular Devices SpectraMax M2) immediately and every minute for 30 minutes to determine optimal pH for enzyme efficiency. Fluorescence at optimal pH was then compared to a commercial β-glucosidase (DesR) as a negative control as above. Thermostability of the putative endo-1,4-β-xylanase was determined by incubating samples containing 0.6 μg of enzyme for 30 minutes at 37°, 50°, 60°, or 70° prior to addition of CMU-X_2_. Samples were then read as before. Control is unincubated endo-1,4-β-xylanase. To calculate kinetic constants, initial rate of product formation (cleavage of CMU from xylobiose) was calculated for 0.6 μg of endo-1,4-β-xylanase at differing concentrations of CMU-X_2_ (10 μM, 15 μM, 20 μM, 40 μM, 100 μM, 150 μM, 200 μM, 250 μM). Kinetic constants *K*_M_ and *k*_cat_ of the putative endo-1,4-β-xylanase were determined using a Lineweaver-Burk plot.

### Plate Assay

BL21 *E. coli* containing pET26b(+)-β-glucosidase, empty pET26b(+) (negative control), or positive control pET28b(+)-DesR [14] were grown overnight in 5 mL LB broth with kanamycin (50ug/ml) at 37°C, shaking. The next day, 1 ml of overnight culture was added to 4 mL of fresh LB and kanamycin. The bacteria were induced with 1mM IPTG for four hours at 37°C, shaking. Then, 1 ml of culture was plated on cellobiose M9 media (33.7 mM Na2HPO4, 22 mM KH2PO4, 8.55mM NaCL, 9.35 mM NH4Cl, 12% cellobiose, 1 mM MgSO4, and 0.03 mM CaCl2) and incubated overnight at 37°C. Colony growth was observed the following day.

## Acknowledgements

We thank Chris Fetter (Dalhousie U.) for guidance in enzyme kinetics calculations. We thank Drs. Steven Hallam and Steve Withers (UBC) for the generous gift of CMU-X_2_ substrate. We thank Zack Armstrong (UBC) for help with troubleshooting enzyme assays and manuscript pre-review. We thank all of the members and mentors of the 2016 and 2017 Dalhousie iGEM teams who launched the porcupine microbiome studies that formed the foundation for the current study.

## Supporting Information

**Table S1.** NCBI Accession Numbers for sequences related to candidate genes from porcupine microbiome from NCBI database

**Figure S1. Alignments** Putative S1A) Endo-1,4-β-xylanase, S1B) β-L-arabinofuranosidase, S1C) β-xylosidase, and S1D) β-glucosidase proteins were aligned to 4 diverse proteins from different bacterial isolates. Alignments were completed using ClustalW, and catalytic sites are annotated with a (*) below to highlight key residues.

**Figure S2. Purification of β-xylosidase** A) Putative β-xylosidase was purified by 6xHIS purification. Red arrow indicates relevant band B) Activity of putative β-xylosidase against *p*-NPX at pH 5.0, and, 6.0 and plotted with positive control; commercial β-xylosidase from *Selenomonas rutinantium* (Megazyme, Ireland).

**Figure S3. Enzymatic activity of endo-1,4-β-xylanase versus unrelated β-glucosidase** Enzymatic activity of endo-1,4-β-xylanase was assessed using cleavage of CMU-X_2_ and emission (450 nm) of CMU at 37°, pH 7, and compared to negative control β-glucosidase DesR.

