## Supplementary Materials for "Discovery and Characterization of Novel Lignocellulose-Degrading Enzymes from the Porcupine Microbiome by Synthetic Metagenomics" (from another version of this preprint: version 1, 2018-03-27)

### RESPONSES TO REVIEWERS

#### Overview:

We previously submitted a PLOS Report to the Breezio platform hosted by PLOS iGEM Collections. In this report we identified a putative β-xylanase enzyme gene from the porcupine microbiome, which was synthesized and partially characterized for xylanase activity. Our study was performed using shotgun metagenomic sequencing from the porcupine microbiome to search for potential lignocellulose-degrading enzymes genes via a metagenomic pipeline. This pipeline allowed the identification of four putative enzyme genes essential for the cellulose- and hemicellulose-degrading pathways. Of the four enzyme genes, β-xylanase was expressed in an *E.coli* model by IPTG induction, however activity could not be detected using a fluorescent assay measuring the cleavage of xylobiose.

In our revised report, we have fully addressed the comments and suggestions put forth by the reviewers. These comments have led to the creation of a new figure (Figure 4), as well as a reorganization of the Interpretation and Future Studies sections. We believe this has strengthened the report, as well as increased clarity. We hope that you and the reviewers will find it suitable for publication in PLOS Channels and Collections Blog.

The major revisions include the following:

1. Figure 2 was deleted as it was not informative to the reader
2. A new figure was created to address Reviewer 3's comments on how our putative β-xylanase homology compares to other characterized β-xylanases.
3. The Interpretation and Future Studies sections were re-organized to improve all around clarity. The Interpretation was re-written to include background on other lignocellulosic biomass degrading sources in the plant and animal kingdoms in response to Reviewer 4. Our plan on incorporating our putative enzymes into a bioreactor model was also added to the Interpretation in response to Reviewers 5 and 6.

#### Detailed responses to comments on "Discovery and Characterization of Novel Lignocellulosic Enzymes from the Porcupine Microbiome"

**Reviewer 1 (lgallagher):**
Hi Dalhousie, Thanks for your submission! Please can you add author affiliations. For an example of how to set these out have a look at University of Calgary's submission here: https://synbioplos.breezio.com/article/4916630916035511997/astroplastic-a-start-to-finish-process-forpolyhydroxybutyrate-production-from-solid-human-waste-using-genetically-engineered-bacteria-to-addressthe-challenges-for-future-manned-mars-missions

**Response:**
**We have addressed the concerns put forth by the review by addition of author affiliations.**

**Reviewer 2 (aaa_2018):**
Do you mean Not Applicable?

**Response:**

**We have addressed the concerns of Reviewer 2 by changing our Ethics Statement from 'Not Available' to 'Not Applicable.'**

**Reviewer 2 (aaa_2018):**
Well done for using protocols.io! I don't see any publicly-visible protocols in your project yet, though.

**Response:**
**We have addressed Reviewer 2's concerns by making all protocols uploaded to protocol.io public.**

**Reviewer 1 (lgallagher):**
It would be great to see a larger version of figures 1 and 2, as the text is a little hard to read at this size.

**Response:**
**Figure 1 was enlarged for the final upload, and Figure 2 was deleted as it was not informative to the reader.**

**Reviewer 3 (TeOH):**
Why didn't you add a tag to your cloned protein in order to be sure of its production but also of it's good functionality, at least in vitro? Moreover, could you add the lower error bar on your figure?

**Response:**
**To answer the reviewer's question, we have designed primers to include the HIS-tag of the pET26b(+) plasmid, however our first experiments used the primer set that added a stop codon before the HIS-tag. This was to allow us to build an operon of multiple enzymes in the degradation pathway of hemicellulose and cellulose. Due to time constraints we never made it to the operon but are currently tagging the putative enzymes in hopes to see our proteins expressed via western blot.**
**In response to the reviewer's concern about Figure 4 (now Figure 3), the error bars were removed entirely as they only represented technical replicates and the authors found it may be misrepresenting the data.**

**Reviewer 3 (TeOH):**
It could be interesting to compare the homology of several other characterized B-xylanase with your putative one and the one of Butyrivibrio sp.

**Response:**
**To address reviewer 3's suggestion, Figure 4 was created to compare the amino acid homology of our putative β-xylanase enzyme against four similar β-xylanase enzymes, including one from the Butyrivibrio spp.**

**Reviewer 4 (Dario):**

Hi, thanks for submitting this interesting study! Although you already explained in the intro the rationale of searching lignocellulosic enzymes specifically in the porcupine microbiome, it might be worth expanding the discussion here on whether other sources (animals, plants, etc.) could be used instead.

**Response:**
**To address reviewer 4's suggestion, paragraph one of the Interpretation was rewritten to include cellulolytic activities of other sources such as fungi, and the microbiomes of ruminids and termites.**

**Reviewer 5 (Serenity):**
What type of system could these enzymes be incorporated into to aid in the production of biofuels?
**Reviewer 6 (nbarvo):**
If similar research is out there, it might be a good idea to discuss biofuel production in your discussion section and how you would incorporate your findings into such a process. Also, you could site previous research to show this is viable and has been done before.

**Response:**
**In response to Reviewers 5 and 6, paragraph 2 of the Interpretation was rewritten to include information on how our expressed β-xylanase will eventually be incorporated into a full operon of genes in the hemicellulose-degradation pathway and expressed in an *E. coli* model as part of a bioreactor process to yield ethanol biofuel.**

**Reviewer 1 (lgallagher):**
Hi Dalhousie, Please clarify why you chose to analyse the gut microbiota of the North American Porcupine, rather than any other animal. Is this due to already available data?
**Reviewer 7 (hw_zt):**
Is there a reason why your team chose to isolate and characterize lignocellulosic enzymes from the porcupine microbiome instead of any other animal? Is this because these enzymes have been found more efficient from previous research?

**Response:**
**In response to Reviewers 1 and 7, our introduction explains that the metagenomic sequencing was obtained in 2016 by the Dalhousie 2016 iGEM team who chose the porcupine as a source for lignocellulose-degrading enzymes as it has a wood-rich diet. We have added some additional description of the findings of the 2016 Dal iGEM team, which was published last year in PLOS ONE (Ref 6).**
