## Supplementary Materials for "Discovery and Characterization of Novel Lignocellulose-Degrading Enzymes from the Porcupine Microbiome by Synthetic Metagenomics" (from another version of this preprint: version 1, 2018-03-27)

Mackenzie Thornbury*, Jacob Sicheri*, Caroline Guinard, David Mahoney, Francis Routledge, Matthew Curry, Mariam Elaghil, Nicholas Boudreau, Angela Tsai, Patrick Slaine, Emma Finlayson-Trick, Landon Getz, Jamie Cook, Dr. John Rohde, Dr. Craig McCormick

*Co-Authors

**Comments**

**lgallagher:** Hi Dalhousie, Thanks for your submission! Please can you add author affiliations. For an example of how to set these out have a look at University of Calgary's submission here: https://synbioplos.breezio.com/article/4916630916035511997/astroplastic-a-start-to-finish-process-for-polyhydroxybutyrate-production-from-solid-human-waste-using-genetically-engineered-bacteria-to-address-the-challenges-for-future-manned-mars-missions

**EmmaDAL:** Mackenzie Thornbury* (1), Jacob Sicheri* (1), Caroline Guinard (1), David Mahoney (1), Francis Routledge (2), Matthew Curry (1), Mariam Elaghil (1), Nicholas Boudreau (1), Angela Tsai (3), Patrick Slaine (1), Emma Finlayson-Trick (1), Landon Getz (1), Jamie Cook (1), Dr. John Rohde (1), Dr. Craig McCormick (1) 1. Department of Microbiology and Immunology, Dalhousie University, Nova Scotia, Canada 2. Department of Neuroscience, Dalhousie University, Nova Scotia, Canada 3. Faculty of Science, Dalhousie University, Nova Scotia, Canada

**lgallagher:** That's great! Thank you.

**Author Contributions**

Conceptualization: The entire team with the help and advice of mentors and supervisors

Parts Design: Jacob Sicheri, Mackenzie Thornbury, Mariam Elaghil; Supervision: Patrick Slaine, Landon Getz,

Bioinformatics: Nicholas Boudreau, Matthew Curry, and Jacob Sicheri; Supervision: Landon Getz

Cloning: Jacob Sicheri, Mackenzie Thornbury, Caroline Guinard; Supervision: Patrick Slaine, Landon Getz, Emma Finlayson-Trick

Expression Assay: Jacob Sicheri, Mackenzie Thornbury; Supervision: Jamie Cook, Landon Getz, Emma Finlayson-Trick

Activity Assay: Jacob Sicheri; Supervision: Jamie Cook, Patrick Slaine

Manuscript Writing: Mackenzie Thornbury, Jacob Sicheri, Matthew Curry, Francis Routledge, Mariam Elaghil, Caroline Guinard, David Mahoney, Emma Finlayson-Trick, Landon Getz

Manuscript Editing: The entire team with the help and advice of mentors and supervisors.

**Abstract**

Plant cell walls are comprised of cellulose, hemicellulose, and lignin, collectively known as lignocellulose. Microorganisms degrade these components to liberate sugars to meet metabolic demands. Using a metagenomic sequencing approach, we previously demonstrated that the microbiome of the North American porcupine (*Erethizon dorsatum*) is replete with novel lignocellulose-degrading enzymes. Here, we report the identification, synthesis and partial characterization of four genes from the porcupine microbiome encoding putative novel lignocellulose-degrading enzymes, including a β-xylanase, endoxylanase, β-glucosidase, and an α-L-arabinofuranosidase. These genes were identified via conserved catalytic domains associated with cellulose and hemicellulose degradation. We cloned the putative β-xylanase into the pET26b(+) plasmid, enabling inducible gene expression in *Escherichia coli* (*E. coli*) and periplasmic localization. We demonstrated IPTG-inducible accumulation of β-xylanase protein, but failed to detect enzyme activity in an assay incorporating a fluorescent xylobiose substrate. Alternative assays may be required to measure activity of this putative β-xylanase. In this report, we describe how a synthetic metagenomic pipeline can be used to identify novel microbial lignocellulose-degrading enzymes, and take initial steps to introduce a hemicellulose-degradation pathway into *E. coli* to enable biofuel production from wood pulp feedstock.

**Financial Disclosure**

Funded by SpringBoard, Dalhousie Medical Research Foundation, Dalhousie University Office of the President, Dalhousie Faculty of Science, Dalhousie Faculty of Medicine, Genome Atlantic, and Dalhousie Department of Microbiology and Immunology.

The funders had no role in study design, data collection and analysis, decision to publish, or preparation of the manuscript

**Competing Interests**

The authors have declared that no competing interests exist.

**Ethics Statement**
Not Available

**Comments**

**aaa_2018:** Do you mean Not Applicable?
**EmmaDAL:** Yes

**Data Availability**
Yes -- all data are fully available without restriction

Metagenomic raw data reads can be found at the NCBI Sequence Read Archive at accession number SRP115632

**Introduction**

The gut microbiota contains thousands of bacterial species, encoding millions of genes (the microbiome), that participate in host physiological processes, such as digestion [1]. The gut microbiome, therefore, is a genetic repository of digestive enzymes. These enzymes, once identified, could be useful in various bioengineering applications, such as biofuel production.

In general, lignocellulose biomass consists of cellulose, hemicellulose, and lignin in a 4:3:3 ratio [2]. Lignocellulose is an attractive substrate for biofuel production via microbial fermentation [3,4]. Prior to fermentation, lignocellulosic biomass must be broken down into its component sugars by enzymatic or chemical degradation. As a hind-gut fermenter, the North American Porcupine, Erethizon dorsatum, has an enlarged cecum packed with microbes to aid in the digestion of lignified plants, coniferous (preferred) and deciduous cambium (inner bark), and flowers [5]. We previously identified that in addition to the common gut microbes, the porcupine microbiome houses specialized lignocellulose-degrading bacteria [6].

Lignocellulosic biofuel production will depend, in part, on the development of synthetic microbial pipelines that effectively degrade the complex polymers found in lignocellulose into simple fermentable sugars. Lignin is believed to be a major hindrance to enzymatic hydrolysis, thus the removal of lignin for biofuel production is crucial. Wi *et al.* recently demonstrated a new pretreatment that improves downstream biocatalytic hydrolysis of lignocellulose and cellulose hydrolysis efficiency [7]. Similar to cellulose, hemicellulose contains potentially useful monosaccharides that can be utilized for ethanol production, such as xylose [8]. Currently, hemicellulose sugars are not widely used for ethanol production, leading to losses in efficiency of biofuel production per input of lignocellulosic biomass.

The primary objective of this study was to first identify enzymes from the porcupine microbiome that have potential to degrade lignocellulose and other polymers found in wood. Next, these proteins need to be expressed to characterize the functionality of the enzymes identified. We generated a synthetic pipeline to *in silico* identify candidate cellulose and hemicellulose degrading enzymes from the metagenomic sequence data from the porcupine, previously published by the 2016 iGEM team [6]. This pipeline identified four unique enzymes, some of which showed highly unique sequences compared to current protein databases. One the identified enzymes, β-xylanase, was successfully cloned and transformed into *Escherichia coli* (*E. coli*) BL21(DE3). Expression was confirmed using an IPTG-induction assay and subsequent analysis of protein production. Next, the activity of the enzyme was measured using a fluorophore-conjugation assay to test enzymatic functionality. The enzyme failed to increase fluorophore activity above background levels. In this iGEM report, we demonstrate successful expression of an uncharacterized β-xylanase identified from the porcupine microbiome and outline the initial steps required to introduce the hemicellulose-degradation pathway in *E. coli*.

**Methods**

All protocols can be found here (https://www.protocols.io/researchers/dalhousie-igem).

**Comments**

**aaa_2018:** Well done for using protocols.io! I don't see any publicly-visible protocols in your project yet, though.
**EmmaDAL:** We will look into this!

*Identification of Open Reading Frames*

Metagenomic analysis was carried out by using previously acquired Illumina MiSeq data [6]. . FASTQC and BowTie2 were used to subsequently inspect reads for overall quality then contaminants from sequencing. Reads were trimmed to 400 bp in length to avoid low-quality sequence ends. MegaHIT alignment software processed reads in FASTq format and assembled reads into longer contigs by identifying overlapping coding regions [9]. Prodigal was used to identify open reading frames (ORFs) by searching sequences in forward and reverse complement [10]. A -c command modifier in Prodigal was used to ensure the program only detected ORFs with both start and stop codons present. Prodigal also used a database of non-canonical start stop codons, as well as ribosome binding sites to identify all ORFs present in sequencing data.

*Predicting Functionality of Putative Proteins*

pHmmer was used to predict the functionality of protein domains [11,12]. Output data from pHmmer was provided in text files. Protein domains and possible functions were identified using the Research Collaboratory for Structural Bioinformatics Protein Data Bank [13]. e-values were calculated to represent the variation between the ORF domain and the domain from the database [13]. Proteins with the lowest e-values were screened against the Basic Local Alignment Search Tool (BLAST) database using pHmmer to identify proteins with major protein domain conservation. Sequences were then optimized for cloning into *E. coli* using IDT's codon optimization tool. ORFs were subsequently synthesized by IDT and used for downstream cloning (Integrated DNA technologies, USA).

*Polymerase Chain Reaction and Gel Extraction*

Putative enzyme genes were PCR-amplified off of the IDT gene blocks using the NEB Phusion High-Fidelity protocol (NEB, M0530S). PCR products were purified using the QIAquick gel extraction kit protocol (Qiagen, 28704) (Table 1).

*Digestion and Ligation*

The target genes and pET26b(+) were digested with the appropriate restriction enzymes for 1 hour at 37 ℃ (NEB) (Table 1). Digested products were subjected to 0.8% agarose gel electrophoresis and purified following the manufacturers protocol (Qiagen, 28704). Following purification, the target gene and plasmid were ligated together using NEB T4 ligase as per manufacturer's protocol (NEB, M0202S). The DNA was quantified using a NanoVue Plus Spectrophotometer (GE Healthcare).

| Enzyme of Interest | Forward Primer | Reverse Primer | Restriction Enzymes Used for Insertion into pET26b(+) (5'-3') |
|---|---|---|---|
| β-glucosidase | GAATTCAAGGA TCCAATGSSCAA GACCCAC | AAAAAACTGCAG CATAGCGGCCGC TGAGCTCTTAAG CTT | SacI/HindIII |
| β-xylanase | AAAAAAGGATC CT ATGGCTAACCC GTACTTACCTG | AAAAAA GAGCTC A TTACTTGGTAAACA CGAACGCC | BamHI/SacI |
| Endo xylanase | GAATTCTGGATCC AATGATCAATAAA TTACTGTCATCGG CATTGGCA | AAAAAACTGCAGCA TAGCGGCCGCTGA GCTCTTAAGCTT | SacI/HindIII |
| Alpha-L-Arabinofuranosidase | GAATTCTGGATCC AATGGGTCTTGCT ATCG | AAAAAACTGCAGCA TAGCGGCCGCTGA GCTCTTCTCGAG | HindIII/XhoI |

**Table 1. PCR primers (forward and reverse) and restriction enzymes sites for the four target enzymes.**

*Transformation into E. coli*

Stbl3 *E.coli* were transformed as previously described with the following exceptions [14]: 5 µL of the ligation was added to 50 µL of *E. coli*, and following heat-shock transformation, 250 µL of LB broth was added during the recovery stage and plated on kanamycin-containing LB plates.The plates were incubated at 37℃ for 18-24 hours. To isolate plasmid DNA, clones were selected and inoculated into 5 mL of LB broth, grown over night and plasmid DNA was extracted by QIAprep spin miniprep kit (Qiagen, 27104). Insertion was confirmed by sanger sequencing (Genewiz).

*Induction Assay*

A single colony of β-xylanase-pET26b(+) transformed into BL21(DE3) *E. coli* was grown in 5 mL LB broth and incubated in a shaking 37˚C incubator at 220 RPM. To induce protein production, 0.1 mM IPTG (Sigma-Aldrich) was added to the culture when an $OD_{600}$ of 0.6 was reached. The culture was incubated at 37°C for 4 hours, at which point the bacteria were lysed with 2x Laemmli buffer to harvest protein for analysis.. Empty vector pET26b(+) (EV) was used as a negative control. Isolated proteins were separated on a 12% SDS-PAGE gel and stained with Coomassie Brilliant Blue (Thermo Scientific) to visualize total protein.

Enzymatic Activity Assay

The enzymatic activity assay was modified from a previously described method [15]. BL21(DE3) *E. coli* were transformed with β-xylanase-pET26b(+) as described previously. At an $OD_{600}$ of 0.6 protein expression was induced using the allolactose mimetic IPTG which triggers transcription from the expression plasmid. Cells were incubated with 0.1 mM IPTG for 4 hours before adding 50 μL of bacteria. Then, 50 μL of bacteria were added to an opaque-walled 96 well plate (Thermo Fisher Scientific). To the bacteria, 50 μL of 200 μM of 4-methylumbelliferyl β-glycosides of xylobiose (CMU-X) in lysis buffer (1% Triton-X100, 50 mM Potassium Acetate at pH 7) was added. The plate was incubated at 37°C for 18 hours, shaking at 220 RPM. After incubation, fluorescence was measured on Tecan Infinite M200 PRO microplate reader with excitation at 365 nm, and emission at 450 nm. Each sample was normalized to LB broth alone. The unconjugated fluorophore, 4-methylumbelliferone, was used as a positive control and uninduced recombinant *E. coli* and pET26b(+) empty vector were used as negative controls.

**Data**

*Mining the Metagenomic Sequencing of the Porcupine Microbiome*

From our pipeline (Figure 1A), four genes, coding for four enzymes involved in cellulose- and hemicellulose-degradation, were identified (Table 2). These hits contained conserved sequences for catalytic domains that are shared with known enzymes sequences (Figure 1B)

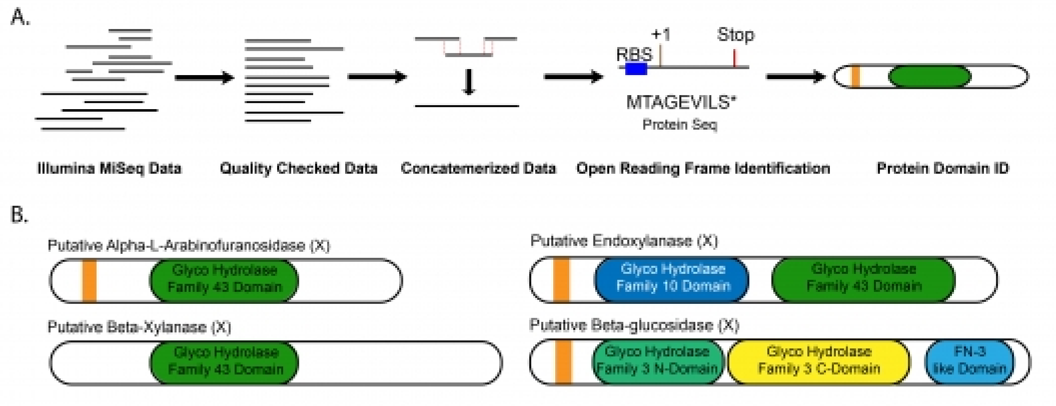


**Comments**

**lgallagher:** It would be great to see a larger version of figures 1 and 2, as the text is a little hard to read at this size.

**Figure 1: Schematic of bioinformatic pipeline and four putative genes of interest.** The bioinformatic pipeline (A) begins with Illumina MiSeq data previously collected [6]. The data is then cleaned up, concatemerized (using MegaHit), open reading frames are identified (using prodigal) and protein sequences of interest are identified by pHMMER and e-value selection. The four protein sequences of interest (B) are shown. The orange bar represents putative localization signals (for import and export to various cell compartments) and various protein domains are indicated.

| Gene (Putative Function) | Function | e-Value |
| --- | --- | --- |
| Beta-Glucosidase | Cleaves beta-1,4 glycosidic bonds between two glucose monomers | 2.1E-125 |
| Beta-Xylanase | Cleaves beta-1,4 bonds between two xylose monomers | 7.6E-157 |
| Endoxylanase | Cleaves internal beta-1,4 bonds in xylose polymers | 8E-94 |
| Alpha-L-Arabinofuranosidase | Cleaves terminal non-reducing alpha-L-arabinofuranoside residues | 3.5E-94 |

**Table 2: Four putative enzyme genes mined from the porcupine microbiome.** Four putative genes with low e-values, indicating high confidence, were chosen to clone into *E. coli*.

*Insertion of β-xylanase into the expression plasmid pET26b(+)*

The four putative enzymes mined from the porcupine microbiome were cloned into the pET26b(+) vector (Figure 1B). pET26b(+) contains an upstream T7 promoter, allowing potent induction of transcription of an inserted ORF after IPTG addition. β-xylanase was the first enzyme successfully cloned by our team, indicated by sanger sequence verification (Figure 2).

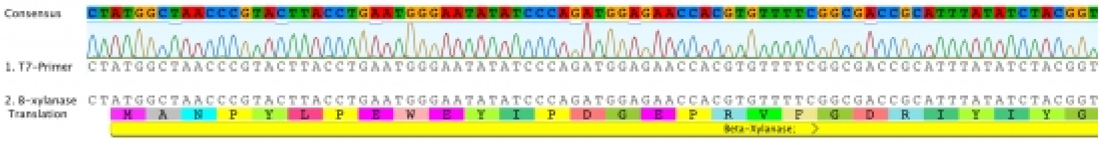


**Figure 2: Cloning of a putative β-xylanase.** A putative β-xylanase identified from the microbiome of the North American Porcupine (Erethizon dorsatum) was synthesized *de novo* by IDT Technologies and cloned into the pET26b expression vector system. The first sequence (T7 primer) indicates the β-xylanase cloned into pET26b(+), the second sequence (β-xylanase Translation) is the original sequence ordered from IDT.

*Candidate β-xylanase Induction Assay*

Recombinant pET26b(+) containing β-xylanase induced by IPTG produced a thick band around 51 kDa, the expected size of our candidate β-xylanase (Figure 3). This band was not observed in lanes containing empty vector (+/- IPTG) and non-induced β-xylanase-pET26b(+). Results were reproduced in technical replicates seen on the left and right side of the marker (Figure 3).

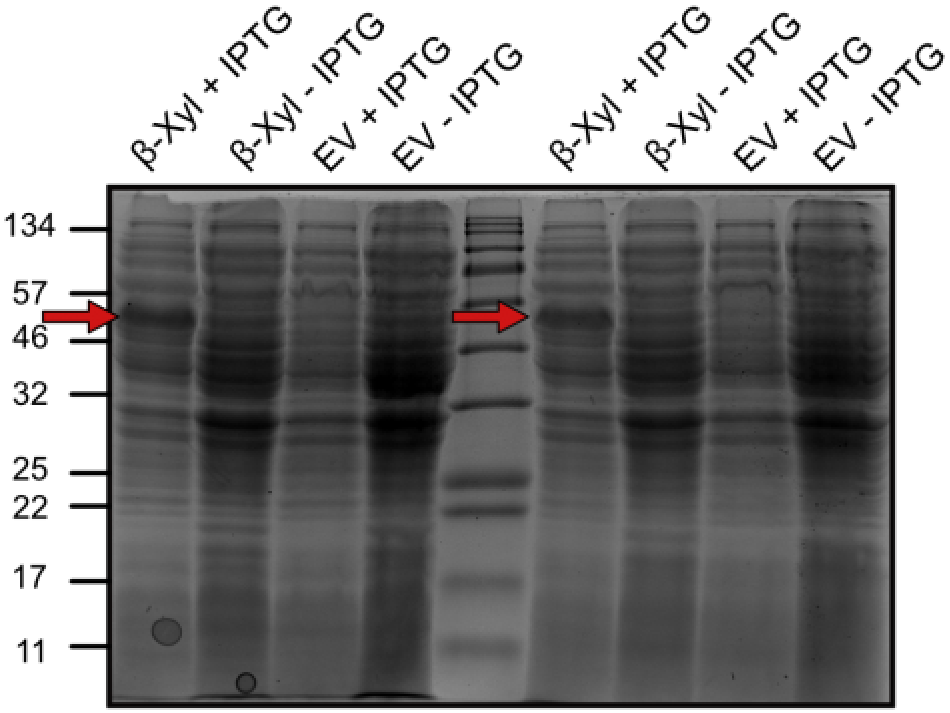


**Figure 3. Expression of β-xylanase in *E. coli*(DE3) via IPTG induction visualized by Coomassie Blue Stain**. Lysates of recombinant or empty vector DNA were induced with IPTG and run via SDS-PAGE, then stained with Coomassie Brilliant Blue. Technical replicates are separated by a molecular weight marker. The red arrows indicate the protein of interest ~51 kDa.

*β-xylanase Enzymatic Activity Assay*

β-xylanase activity was tested using a technique developed by the Hallam Lab [15]. The assay requires the use of fluorophores conjugated to a xylose sugar via a β-1,4 glycosidic bond. If β-xylanase was active, it would cleave the bond resulting in measurable fluorescence at 450 nm. The recombinant β-xylanase was induced as previously described and fluorescence was measured. There was no significant difference between the induced β-xylanase when compared to the uninduced β-xylanase and empty vector control. Fluorescence was ~1.5 logs greater in the unconjugated fluorophore sample in comparison to the β-xylanase sample (Figure 4).

**Comments**

**TeoH:** Why didn't you add a tag to your cloned protein in order to be sure of its production but also of it's good functionality, at least in vitro ? Moreover, could you add the lower error bar on your figure ?
**MackenzieRT:** Hi Teoh, We have designed primers to include the his-tag of the pET26b(+) plasmid, however our first experiments used the primer set that added a stop codon before the his tag. This was to allow us to build an operon of enzymes once cloning was successful. Due to time constraints we never made it to the operon, but are currently tagging the putative enzymes in hopes to see our 'proteins' expressed via western blot. We will look into adding a lower error bar.

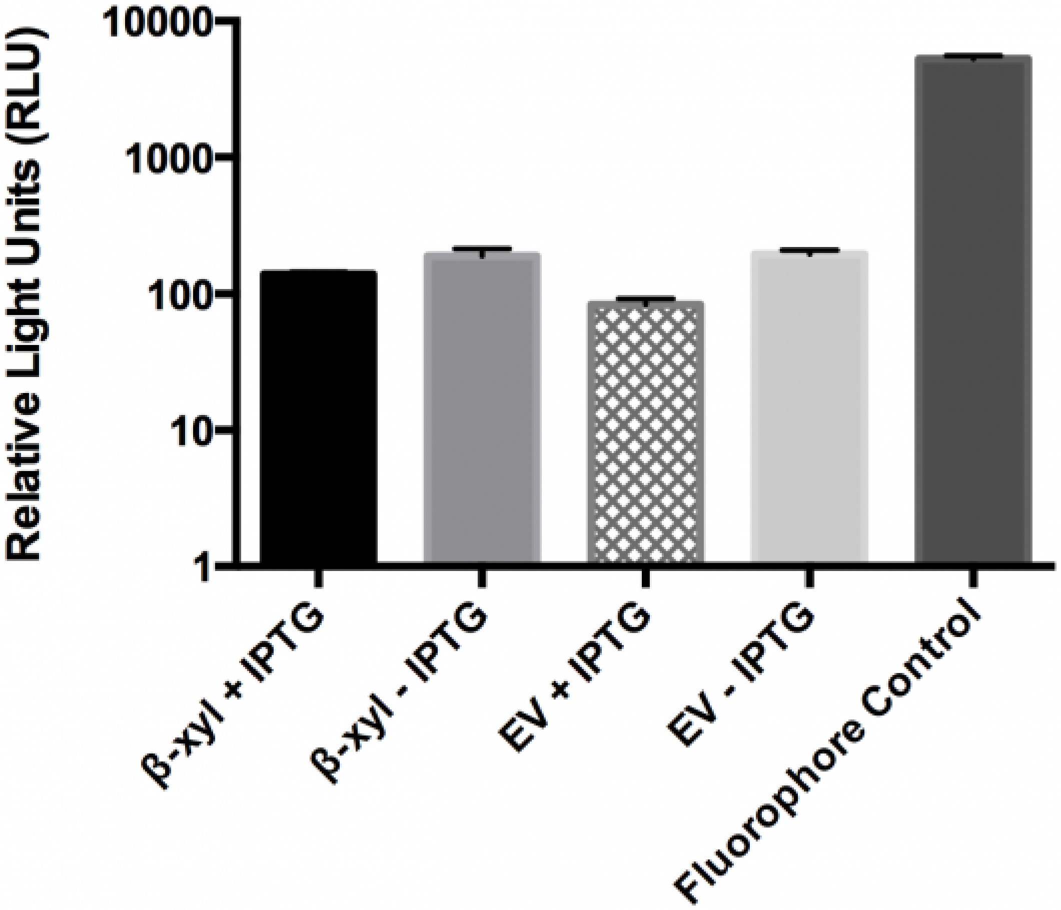


**Figure 4. Analysis of enzymatic activity via a fluorophore cleavage assay.** Following IPTG induction, the transformed *E. coli* were lysed and mixed with a solution containing the xylose conjugated fluorophore. After 18 hours, relative light units (RLU) were measured on a spectrophotometer and graphed via Prism 6 (Graphpad Inc.) Error bars represent the standard deviation from 3 technical replicates.

**Interpretation**

Mining the porcupine gut microbiome resulted in the identification of four putative enzymes involved in cellulose and hemicellulose-degradation. These enzymes are unique, but contain the conserved activity domains of previously characterized enzymes of the same type. Of these four candidate enzymes, we investigate an uncharacterized β-xylanase enzyme with 75% homology to a *Butyrivibrio sp.* β-xylanase outlined on the NCBI protein database (CDC35707.1). Through the metagenomic pipeline, we were able to screen for protein activity with ease, limited only by the time constraint of the study. The metagenomic sequencing from the porcupine microbiome can be screened for other specialized enzymes, but due to the specific hypothesis of this study, only hemicellulolytic and cellulolytic enzymes were pursued. This study lays the foundation for future iGEM teams to use *in silico* pipelines to identify novel enzymes from metagenomic sequencing as a resource for industrial platforms.

**Comments**

**TeoH:** It could be interesting to compare the homology of several other characterized B-xylanase with your putative one and the one of Butyrivibrio sp.
**EmmaDAL:** Definitely something we could look into doing! Thank you TeoH

The putative β-xylanase appeared in our Coomassie Brilliant Blue stain of SDS-PAGE (Figure 2), confirming the overexpression of the protein in this system. There was no signal above background in the functional fluorochrome assay because our protein was unable to cleave the substrate to release the fluorophore (Figure 3). This could be attributed to environmental conditions, such as oxygen and pH levels in the porcupine stomach that were not reflected in our experiment. The experiment was conducted in an aerobic environment while the porcupine hindgut is an anaerobic environment [16]. Oxygen may be an inhibiting factor in the function of β-xylanase. It is possible that the putative β-xylanase is sensitive to oxygen, specifically the reactive oxygen species that are generated in oxygen-rich environments. This has been shown before in nitrogenase enzymes found in anaerobic bacteria such as *Azotobacter* and *Cyanothece* species [17,18]. Another possible explanation could be that the candidate β-xylanase is designed to function at a pH higher or lower than pH 7.0, the pH at which the fluorophore assay was done. The pH of the environment impacts protein folding, stability and function, a proteins optimal pH is dependent on the pKa values of it’s amino acid composition [19,20].This is relevant to our project as the porcupine’s digestive tract fluctuates based on region, but can reach a pH as low at pH 4.5 which was not reflecte din our original experimental design [21].

**Future Studies**

*New Methods of Enzyme Discovery*

An alternative way to identify truly novel enzymes would be to build a functional metagenomic library. Our current method focused on comparing the metagenomic sequencing of the porcupine microbiome to previously identified enzyme gene sequences. This method limits enzyme discovery, and every new enzyme hit had the possibility of being inactive as it is difficult to predict function without experimental evidence. Functional metagenomics, such as through a cosmid or F-plasmid systems, rely completely on functional screens to discover hits. This way, the function is known before the sequence. This can lead to discovery of new types of enzymes, that may have little to no homology to previously discovered enzyme species. Recently, Cheng *et al*. discovered three novel β-galactosidase enzymes, two of which had conserved domains, and one of which was part of a previously undiscovered enzyme family [22]. While less high-throughput, we believe creating a functional metagenomic library from the porcupine microbiome DNA would be a viable way to continue to search for enzymes in the cellulose and hemicellulose degradation pathways.

**Comments**

**Dario:** Hi, thanks for submitting this interesting study! Although you already explained in the intro the rationale of searching lignocellulosic enzymes specifically in the porcupine microbiome, it might be worth expanding the discussion here on whether other sources (animals, plants, etc.) could be used instead.
**MackenzieRT:** Thanks Dario, that's a good direction for our discussion!

*Alternative Enzyme Activity Assays*

Going forward, several other assays can be performed to assess the enzymatic activity of the enzymes produced by transformed *E. coli.* One such assay to measure the effectiveness of β-xylanase is the Somogyi-Nelson method based upon measuring the reducing power of sugars in solution [23]. The Somogyi-Nelson method relies on cupric ion reduction by a reducing sugar such as glucose and xylose [23]. Therefore, this method will also measure all reducing sugars from the growth media. To account for this, growth media will have to be prepared containing alternate carbon sources. By measuring xylose monomers produced by the β-xylanase reaction with it's substrate xylan, we can calculate β-xylanase activity.

Another assay suited to measure endo-glucanase activity is the agar media based Congo Red staining assay. As pH decreases the colour of Congo Red darkens, this allows for measurement of the reaction between carboxymethylcellulose and endoglucanase which yields a cellobiose molecule as well as a phosphate group [24]. The phosphate group causes a lowering of pH in that specific location in the media allowing visualization of enzymatic activity qualitatively [24]. However, for this assay to be effective, glucose must not be used as the growth media’s carbon source. Since *E. coli* metabolic pathways for glucose lead to a decrease in pH, a solution to this issue is to use xylose (a sugar *E. coli* can metabolize naturally) as a carbon source which does not lead to a pH decrease upon metabolism [25]. For more quantitative analysis of carbohydrate content, which would allow precise calculation of enzyme activities, samples may be prepared for high performance liquid chromatography using a column designed for carbohydrate analysis [26]. This would be most effectively done in collaboration with a laboratory where HPLC is routine.

    (https://www.google.com/url?q=http://www.rcsb.org&sa=D&ust=1518407432561000&usg=AFQjCNGdhIO82Dgtm5fvhx3iO7K9Z8ifCw)
14. (https://www.google.com/url?q=http://www.rcsb.org&sa=D&ust=1518407432561000&usg=AFQjCNGdhIO82Dgtm5fvhx3iO7K9Z8ifCw)Froger

A, Hall JE. Transformation of plasmid DNA into *E. coli* using the heat shock method. J Vis Exp. 2007 Aug 1;(6):253. doi:10.3791/253.
15. Chen HM, Armstrong Z, Hallam SJ, Withers SG. Synthesis and evaluation of a series of 6-chloro-4-methylumbelliferyl glycosides as fluorogenic reagents for screening metagenomic libraries for glycosidase activity. Carbohydr Res. 2016 Feb 8;421:33-39
16. Karasov WH, Douglas AE. Comparative digestive physiology. Compr Physiol. 2013 Apr;3(2): 741-783
17. Zhang X, Sherman DM, Sherman LA. The uptake hydrogenase in the unicellular diazotrophic cyanobacterium *Cyanothece sp*. strain PCC 7822 protects nitrogenase from oxygen toxicity. J Bacteriol. 2014 Feb;196(4):840-849.
18. Dilworth MJ, Kennedy IR. Oxygen inhibition in *Azotobacter vinelandii* Some enzymes concerned in acetate metabolism. Biochim Biophys Acta. 1963;67: 240-253.
19. Alexov E. Numerical calculations of the pH of maximal protein stability. FEBS J. 2004 Jan;271(1):173-185
20. Pace CN, Grimsley GR, Sholtz JM. Protein Ionizable Groups: p*K* Values and Their Contribution to Protein Stability and Solubility. J Biol Chem. 2009 May; 284:13285-13289.
21. Vispo C, Hume ID. The digestive tract and digestive function in the North American porcupine and beaver. Can J Zool. 1995; 73(5): 967-974
22. Cheng J, Romantsov T, Engel K, Doxey AC, Rose DR, Neufield JD, Chares TC. Functional metagenomics reveals novel β-galactosidases not predictable from gene sequences.PLoS ONE. 2017;12(3): e0172545.
23. Nelson N. A photometric adaptation of the Somogyi method for the determination of glucose. J biol Chem. 1944 May 1;153(2):375-80.
24. Meddeb-Mouelhi F, Moisan JK, Beauregard M. A comparison of plate assay methods for detecting extracellular cellulase and xylanase activity. Enzyme Microb Technol. 2014 Nov 1;66:16-9.
25. Han K, Lim HC, Hong J. Acetic acid formation in *Escherichia coli* fermentation. Biotechnol Bioeng. 1992 Mar 15;39(6):663-71.
26. Honda S. High-performance liquid chromatography of mono-and oligosaccharides. Anal Biochem. 1984 Jul 1;140(1):1-47.

**Comments**

**lgallagher:** Hi Dalhousie, Please clarify why you chose to analyse the gut microbiota of the North American Porcupine, rather than any other animal. Is this due to already available data?

> **EmmaDAL:** Hello Larch, this years iGEM project is a continuation of our project last year when we were interested in determining if the porcupine microbiome, due to its diet, could be mined for certain enzymes, such as cellulases. From that project, we were able to conclude that the porcupine microbiome could indeed be used as a mine for enzymes involved in lignocellulose degradation (as could the beaver). This years project is a continuation, and so we focused on specific enzymes involved in lignocellulose degradation. Does that help clarify our rationale, or would you like me to go more in depth?
>
> **MackenzieRT:** The availability of samples also played a roll in this decision. As porcupines only live on land, their fecal samples are easy to collect. Beavers spend a significant amount of time in the water and their fecal samples are often contaminated with water dwelling microbes.

**hw_zf:** Is there a reason why your team chose to isolate and characterize lignocellulosic enzymes from the porcupine microbiome instead of any other animal? Is this because these enzymes have been found more efficient from previous research?

**Serenity:** What type of system could these enzymes be incorporated into to aid in the production of biofuels?

**MackenzieRT:** Hi Serenity, Dalhousie iGEM looked into two main models, both E. coli and S. cerevisiae. Our 2016 team did preliminary studies that showed yeast were susceptible to turpenes that would be found in softwood lumber feedstock, so we decided to go with a duel model. Our model started with genetically engineered E. coli with our uncharacterized enzymes to produce the sugar that S. cerevisiae could metabolize into ethanol biofuel. Let me know if you'd like me to expand further.

**nbarvo:** If similar research is out there, it might be a good idea to discuss biofuel production in your discussion section and how you would incorporate your findings into such a process. Also, you could site previous research to show this is viable and has been done before.
